# Two newly cultivated eukaryotrophic flagellates represent distinct anaerobic lineages within Rhizaria

**DOI:** 10.1101/2024.09.01.610664

**Authors:** Yana Eglit, Maggie Lawton, Alastair G.B. Simpson, Ryan M.R. Gawryluk

## Abstract

Endomyxans are a poorly sampled and incompletely resolved aggregate of Rhizarian lineages that fall outside Filosa and Retaria. Among them, “Novel Clade 12” (NC12; Bass et al. 2009) is an environmental clade comprised primarily of sequences derived from anoxic sediments, hitherto lacking a morphologically-characterised representative. We have cultivated a marine anaerobic eukaryotroph, SSF, that we identify as the first representative of NC12. SSF is a teardrop-shaped cell with two unequal flagella emerging a third of the way down the cell behind a distinctive row of refractile globules. The posterior end of the cell is filled with food vacuoles. There is a surface thickening discernible in light microscopy. We also describe another distinct anaerobe eukaryotrophic lineage, also cultivated from marine sediment: PG. It consists of large pyriform cells with a substantial trailing “tail” and two unequal flagella, the posterior exceptionally long. In small subunit ribosomal RNA gene phylogenies, it falls outside the characterised clades and forms a distinct novel rhizarian lineage in its own right. Together, SSF and PG represent two additional independent adaptations to anoxic conditions within Rhizaria.

## Introduction

The tree of eukaryotes is composed of a number of supra-kingdom level clades, or supergroups (Burki et al., 2020). One such group, the Sar (Stramenopiles, Alveolates, and Rhizaria), is particularly vast and includes familiar organisms like ciliates and diatoms within alveolate and stramenopile subclades, respectively. Comparatively less well-studied are members of the third subclade, the rhizarians; this group encompasses a bewildering diversity of cell morphologies and lifestyles (Burki and Keeling, 2014). Within it, Retaria is comprised of exceptionally complex giant amoebae foraminifera and polycystine radiolarians (Suzuki and Not, 2015), whereas Filosa is a robust clade dominated by amoeboflagellates but also photosynthetic amoebae (Archibald, 2015; Macorano and Nowack, 2021) and even an osmotroph (Feng et al., 2021).

The rest of rhizarian diversity currently falls within “Endomyxa”, a loose assemblage of poorly resolved lineages (Krabberød et al., 2017) including eukaryotrophic vampyrellid amoebae, large-celled gromiids, parasitic phytomyxids and ascetosporea, and the reticulated amoeba *Filoreta*. Endomyxa are also host to a number of environmental clades identified over the past three decades: most notably “Novel Clades” 10-12. Novel Clade 10 (NC10) and Novel Clade 11 (NC11) are freshwater and contain raptorial eukaryotrophs *Aquavolon* (Bass et al., 2018) and *Lapot* (Irwin et al., 2019), and *Tremula longifila* (Howe et al., 2011), respectively. Novel Clade 12 (NC12), on the other hand, is largely composed of sequences derived from marine anoxic sediments and currently lacks morphological representation altogether. Better sampling of Endomyxans, especially of the Novel Clades and other current environmental lineages, would be integral not only to resolving the branching order of Endomyxa, but also pinpointing the Rhizarian root.

Numerous lineages of eukaryotes have independently adapted to anaerobic environments. While most research efforts have been focused on medically- and economically-important parasitic groups (e.g. *Giardia* or *Trichomonas*), the majority of anaerobes are free-living heterotrophs and considerably less well-studied. They are distributed across the tree of eukaryotes, and feature varying degrees of independently reduced mitochondria termed mitochondrion-related organelles (MROs) (Leger et al., 2019; Roger et al., 2017). While each independent anaerobic lineage features its own unique adaptations and modifications, there are parallels—in terms of gene loss, retention, and gain in diverse anaerobic lineages—that lead to hypotheses of a generalised sequence of events that accompany evolution under anoxia (Stairs et al., 2015). For example, electron transport chain complexes III and IV, involved in the reduction of molecular oxygen, and lost earlier than complexes I and II, which play important roles in recycling of reduced cofactors (Gawryluk and Stairs, 2021). In two cases thus far, the MRO has been lost altogether following the acquisition of a cytosolic alternative to typically-mitochondrial iron-sulfur cluster biogenesis in late stages of anaerobic adaptation (Karnkowska et al., 2016; Williams et al., 2024). Each newly discovered independent anaerobic lineage adds further insight into the process.

Among Rhizaria, *Brevimastigomonas motovehiculus* and mikrocytids (*Mikrocytos mackini* and *Paramikrocytos* spp.) are the two independent lineages of anaerobes with reduced MROs characterised thus far. *Brevimastigomonas* is a free-living bacterivorous filosan that thrives in anoxia, but also tolerates oxic conditions (Gawryluk et al., 2016) whereas *Mikrocytos* and *Paramikrocytos* are endomyxan parasites of shellfish and arthropods (Burki et al., 2013; Onuţ-Brännström et al., 2023). Among foraminifera, *Globobulimina* contains a denitrification pathway (Woehle et al., 2018) but no published evidence yet shows mitochondrial reduction. Predominantly sampled from anoxic sediments, NC12 is a likely (though not definitive) candidate for an additional independently-derived anaerobic lineage of rhizaria.

Free-living protists that feed upon other eukaryotes, or eukaryotrophs, present a particularly understudied ecological niche—owing partly to the difficulty in establishing stable cultures (Tikhonenkov, 2020). Most of the recent super-group-level novel lineage additions to the tree of eukaryotes have been eukaryotrophs (Janouškovec et al., 2017; Lax et al., 2018; Tikhonenkov et al., 2022), even among the previously exclusively algal archaeplastids (Gawryluk et al., 2019; Schön et al., 2021). Eukaryotrophy among anaerobic flagellates has not only been rarely studied, but assumed to be extremely rare based on the much lower proportion of eukaryote biomass detected in anoxic samples; it was previously proposed to be perhaps bioenergetically unlikely in those conditions (Fenchel, 2012). Thus, one would expect that searching for anaerobic eukaryotrophs is a productive approach for characterising significant novel lineages, including some of the environmental clades.

Here, we present isolates of two distinct anaerobic eukaryotrophic lineages within Rhizaria that we have successfully cultivated using breviates as prey. Isolates PCE-SSF and RB2-SSF fall within NC12 and thus the first characterised representative of that clade. PG, on the other hand, represents a novel distinct rhizarian lineage in its own right, previously undetected by environmental sequencing methods.

## Materials & Methods

### Organisms

All isolates were derived from enrichment in 3% LB (final concentration; v:v) in autoclaved natural seawater from different sites in Canada. Locales QSI, TBB1, Saa, and RB2 are tidal mud flats on the Pacific coast of Canada, whereas PCE is a subtidal eelgrass meadow in Cavendish, PEI on the Atlantic coast (see Supplemental Table 1—Sampling Location Table—for sampling details). To establish di-eukaryotic cultures, one to three predator cells were manually picked with a fine glass capillary into cultures of undescribed breviates (Breviata, Obazoa) that they were empirically determined to feed on. All cultures were maintained at room temperature by weekly transfers of a 1 mL innoculum into a week-old culture of the respective prey breviate culture in 15mL conical tubes filled to 12mL with 3%LB in autoclaved natural seawater derived from Halifax, NS, Canada (Northwest Arm) or 4%LB in seawater derived from Victoria, BC, Canada (Cadboro Bay). Prey breviates were grown separately to maximize food availability for the predators.

### Light microscopy

Light microscopy was done using a Nikon Eclipse Ti-2 microscope with a Nikon Digital Sight 10 camera, or a Zeiss Axiovert 200M microscope, with an AxioCam M5 camera. For morphological documentation, cells were imaged in chamber slides made by sealing the edges of a coverslip with vaseline; these chamber slides were allowed to incubate for a minimum of 12h, usually 24-48h, to allow the cells to acclimatise to the glass surfaces. For cell size measurements, aliquots of cultures of PCE-SSF, RB2-SSF, and QSI-PG at 4 days post transfer were transferred to vaseline-sealed chamber slides, incubated for 28-30h, then imaged under 20x phase contrast with the 1.5x tube lens in. Analyses were done using FIJI v.1.53c (Rasband, 1997; Schneider et al., 2012).

### SSU rRNA gene sequencing

To determine the sequences of the various target SSU rRNA genes, we first amplified genomic DNA by whole genome amplification. Briefly, a single cell was picked from each respective di-eukaryotic culture, washed in filtered medium, lysed by 4-6 rounds of freeze-thaw using liquid nitrogen, and amplified by Multiple Displacement Amplification (MDA) at 30°C for 2h using the Illustra GenomiPhi v3 kit (GE Healthcare). The SSU rRNA gene from each isolate was then amplified by PCR using the MDA product as template. PCR conditions and primer sets used for each isolate are described in Supplemental Table 2. Amplicons were separated by electrophoresis through a 1% (w:v) agarose gel, then purified by gel extraction (QIAquick Gel Extraction kit; cat#28115). Purified amplicons were Sanger-sequenced by Génome Québec or by Eurofins Genomics and sequences assembled in Geneious R10 (Kearse et al. 2012).

In the cases of isolates PCE-SSF and QSI-PG, SSU rRNA gene sequences were confirmed and/or improved with Illumina NovaSeq PE150 reads. Here, individual cells were manually picked, as above, and genomic DNA was amplified using the 4BB TruePrime Whole Genome Amplification (WGA) Kit (4basebio, Madrid, Spain), with a 6 hour incubation at 30ºC. Sequences were checked for quality with fastqc (v0.12.0) and normalized with bbmap (v38.86). Adapters and poly-G sequences were trimmed with fastp (v0.23.4). Sequences were assembled with Spades (v3.15.4), using the single-cell assembly mode. Full ribosomal genes were extracted from assemblies using barrnap (v0.9).

### Phylogenetics

A phylogenetically broad collection of rhizarian SSU rRNA gene sequences, along with stramenopiles and alveolates as outgroups, was expanded by a profile alignment of the newly generated sequences using MUSCLE (Edgar, 2004) and manually curated in SeaView v. 5.0.4 (Gouy et al., 2010), then trimmed with gblocks (Castresana, 2000) and manually edited. Maximum likelihood phylogenetic trees were reconstructed from the trimmed alignment (1298 sites and 105 taxa) with IQ-TREE multicore version 2.3.1 COVID-edition (Minh et al., 2020; Nguyen et al., 2015) under the GTR+Γ+I model with support values determined by 200 non-parametric bootstrap replicates.

## Results

We established stable di-eukaryotic cultures of three isolates of “SSF” (Saa-SSF, PCE-SSF, and RB2-SSF) and two isolates of “PG” (TBB1-PG and QSI-PG). All isolates were maintained on a cultured undescribed breviate PCE-brev except Saa-SSF—grown on an undescribed breviate Saa-brev 100% identical to PCE-brev by SSU rRNA gene sequence—and TBB1-PG, which was grown on a larger undescribed breviate LRM1b. Attempts to grow TBB1-PG on PCE-brev and QSI-PG on LRM1b failed. Both SSF and PG could not be grown in absense of the breviate, nor would they tolerate oxygen exposure, even though in the case of TBB1-PG the breviate itself was surprisingly tolerant of oxygen. Lastly, when exposed to air in droplets on slides, both SSF and PG would round up and die similarly to other anaerobic free-living flagellates. This points to SSF and PG being both eukaryotrophic and anaerobic.

### SSF morphology

SSF has a teardrop-shaped cell ∼10-25 μm long with two unequal flagella emerging halfway or two-thirds of the way up the cell (Fig 1). It is strikingly plastic and can be distorted with large quantities of phagocytic vacuoles, and squirms with its whole cell body along with flagellar waves. The cell glides passively upon a long posterior flagellum, the distal end of which is usually attached to the surface. The anterior third of the posterior flagellum is involved in side-to-side wave movement. The proximal end of anterior flagellum is attached to the cell: this attachment ends in a stiff cytoplasmic rostrum (Fig 1 A,D,F). Beneath or often to the left of the flagellum is a distinctive row of conspicuous refractile globules (Fig 1B-D, F). The anterior and posterior flagella each emerge from a corresponding ventral depression, the sides of which are delineated by stiff cytoskeletal structures (Fig 1A, G). Immediately next to the flagellar insertion site is the nucleus (Fig 1D). Phagocytic vacuoles, often abundant, are localised in the posterior end of the cell (Fig 1B, C, E). A thickening discernible under light microscopy is associated with the surface of the cell (Fig 1H). Dispersed throughout the surface are numerous small granules presumed to be extrusomes (Fig 1H, G). Cells can be found with trailing strands of cytoplasm with a granular surface, gradually retracted (Fig 1L). Swimming cells have a snakey, ‘squishy’ movement.

**Figure 1.**
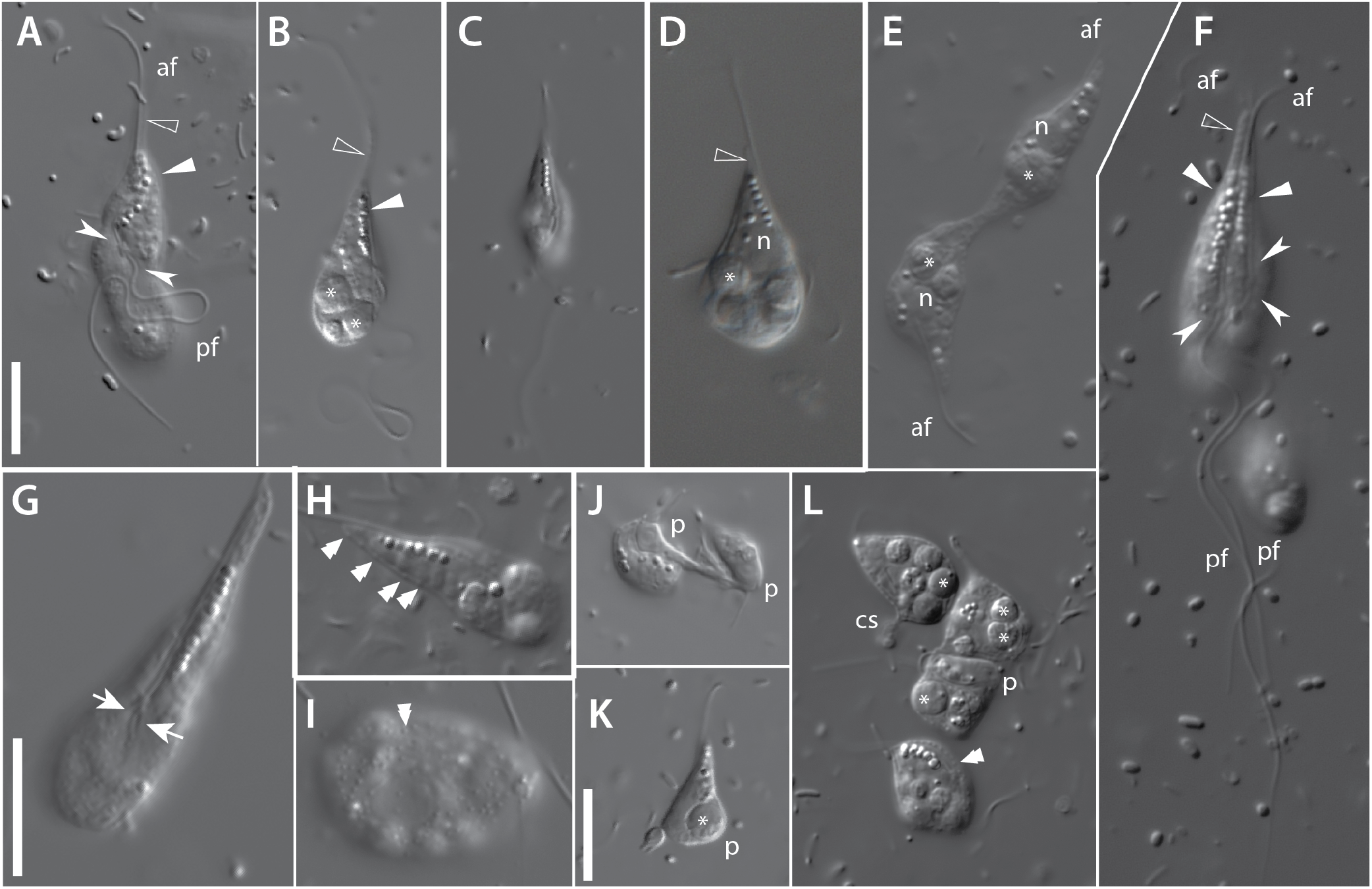
General morphology of SSF isolates PCE-SSF (**A-B, E-G, I-L**), RB2-SSF (**C, H**), and Saa-SSF (**D**) inferred by light microscopy using DIC optics. **A-B**) Ventral views of a slightly flattened specimen (**A**) and a more typically posed cell (**B**), revealing the characteristic row of globules (solid arrowheads), and ventral grooves containing flagellar insertion sites (barbed arrows) for the anterior and posterior flagella (af and pf, respectively). Along the anterior flagellum is a thin cytoplasmic “rostrum” (hollow arrowhead). Numerous phagocytic vacuoles (asterisk) are visible in the posterior end of the cell. **C**) A smaller individual of isolate RB2-SSF. **D**) Lateral optical section through an individual of isolate Saa-SSF additionally showing the nucleus (n). **E-F**) Dividing cells of isolates PCE-SSF. The cells in (**E**) are in late division, whereas the double-rostrumed individual (**F**) is presumably in earlier stages. **G**) Detail of the posterior ventral groove, including edge structures (arrows). **H-I**) Detail of surface granules (double arrowheads), presumably extrusomes, traversing a conspicuous membrane thickening (**H**) and abundantly dispersed throughout the cell surface in an irregular fashion (**I**). **J-L**) Scenes of predation by isolate PCE-SSF upon the breviate prey (p). Early predation (**J**) and a completely engulfed cell (**K**). Several individuals can be seen attempting to engulf a single prey cell (**L**). Note the row of granules visible along the edge of the anterior ventral depression. Sometimes cells form a temporary trailing cytoplasmic strand (cs). Scale bar in **A** and **K** applies to **A-F** and **J-L**, scale bar in **G** applies to **G-I**. All scale bars are 10 μm.

### SSF feeding and division

SSF has been documented feeding on its breviate prey (Fig 1 J-L). Following initial contact with prey, SSF usually continues past. The breviates appear increasingly tattered and eventually round upon. Upon contact with a detached or partly rounded up cell, SSF (not necessarily the same cell as in the initial encounter) contacts it again and then phagocytoses it (Fig 1J, K). Several SSF cells can attempt to ingest a single prey cell (Fig 1L). Prey ingestion occurs at any site on the cell surface, often far from the row of refractile globules (Fig 2J).

**Figure 2.**
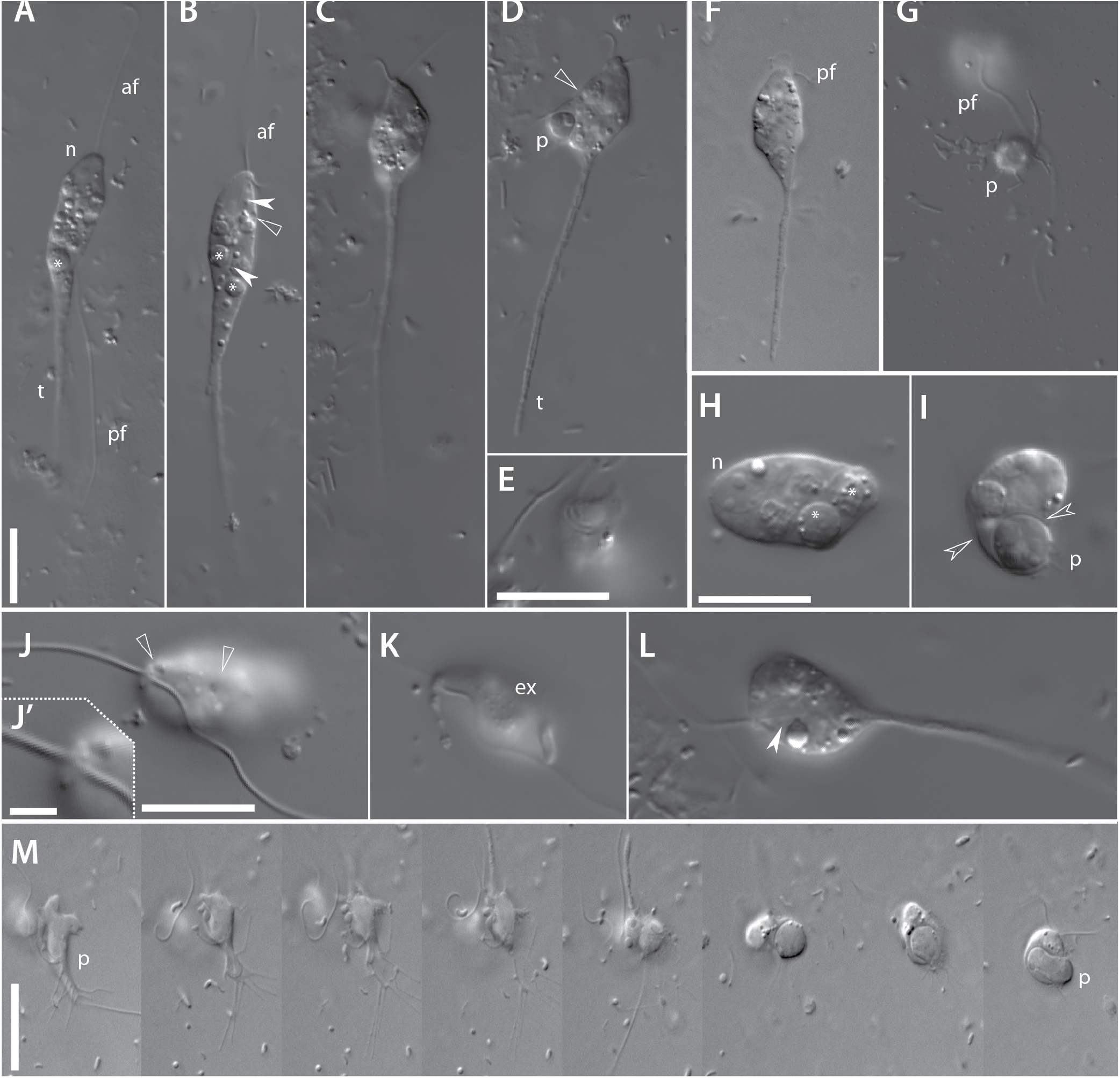
General morphology of PG isolates QSI-PG (all except **F** and **M**) and TBB1-PG (**F, M**) as inferred by light microscopy using DIC optics. **A-D**) General views of QSI-PG. In many cases, both anterior (af) and posterior (pf) flagella can be seen emerging from the anterior end of the cell, as well as a long cytoplasmic ‘tail’ (t). Phagocytic vacuoles (asterisks) can be abundant throughout the cell, as well as an anterior vesicular nucleus (n). An internal rod (barbed solid arrowheads) can be seen in the middle of the cell in (**B**). Small vesicles (hollow arrowhead) are localised at the periphery. A prey cell (breviate PCE-brev; p) can be seen phagocytosed (**D**). **E**) Close up of a flagellum pressed against the cell surface, forming a transient “channel”. **F**) Whole cell view of isolate TBB1-PG. **G**) A prey breviate (p) cell recently contacted by QSI-PG at the top of the frame showing disintegrating filopodia. **H**) Close up of optical section through the cell showing the nucleus, as well as digestive vacuoles in different stages of digestion. **I**) Another optical section through the cell showing an outer layer just discernible by light microscopy (hollow barbed arrowheads). This layer also surrounds the freshly phagocytosed prey. **J**) A close up of the surface of the cell at the enterior end showing flagellar insertion (inset **J’**) as well as two peripheral vesicles. **K**) Another view of the surface cell showing small granules, presumably extrusomes (ex). **L**) Optical section through a cell showing showing the central internal rod. **M**) Time series of a TBB1-PG cell contacting, disrupting, and phagocytosing its breviate prey (isolate LRM1b). Scale bar in A applies to **A-D** and **F-G**; scale bars in **E, H**, and **J** apply to **E** and **H-L**. All scale bars are 10 μm except for **J’**, where it is 2 μm.

Cells with two sets of flagella, oriented in parallel, are relatively common in well-fed cultures (Fig 1F), implying SSF may spend considerable time in that stage. These cells have up to two rostrums and, notably, duplicated anterior rows of globules, usually to the left of each flagellum (Fig 1F). Cells are capable of feeding in this stage. They can have one nucleus or two, presumably indicating different stages of division. Other cells can be found still connected by a cytoplasmic strand (Fig 1E).

### PG morphology

PG (Fig 2) has a pear-shaped cell ∼15-25 μm with a long distinctive trailing “tail” of cytoplasm, up to four cell lengths long, pointing slightly upwards as the organism glides upon its exceptionally long posterior flagellum (Fig 2A-D, F). Together with the “tail”, some cells can be well over 50 μm long. Two unequal flagella emerge from the anterior tip of the cell (Fig 2B-C, J), with the posterior one quite long (Fig 2A). The organism glides passively upon its posterior flagellum. The nucleus is located at the anterior end of the cell by the flagellar insertion site (Fig 2A). Phagocytic vacuoles, sometimes numerous, can be found throughout the rest of the cell (Fig 2A-C, H). A stiff internal rod can be seen inside the cell body proper (Fig 2B, L). The cell is highly plastic and its shape changes depending on cell movement and feeding status. When pressed against the cell surface, the flagellum forms a transient channel (Fig 2E).

While small refractile globules are present throughout the cell, they do not present in rows or any other particular arrangement. The cell surface has an associated thickening discernible by light microscopy (Fig 2I), that is also visible around freshly phagocytosed cells. Immediately underneath the cell surface are small vesicles that may grow and shrink over several minutes (Fig 2B, J). The surface itself is covered by a dispersion of fine granules, presumed to be extrusomes (Fig 2K). This granulation continues into the cytoplasmic “tail”. Occasionally, trailing cytoplasmic strands may form transiently and them be retracted (Fig 2K).

### PG feeding

PG is a voracious predator of its breviate prey. Upon surface contact (of any part of the cell) with a breviate, the prey filopodia rapidly disintegrate over several seconds (Fig 2G, M). During this time, PG may move away. Dead and dying prey cells are then phagocytosed (Fig 2D, I, M), sometimes several at a time. As with SSF, predation appears to occur in two decoupled steps, although PG may return and devour its prey only seconds following initial contact.

## Phylogenetics

The SSU rRNA gene phylogeny recovered both Filosa and Retaria with high support, as well as the phytomyxids, vampyrellids, gromiids, and ascetosporeans (Fig. 3). Environmental clades ‘Endo4’ and ‘Endo5’ were recovered with high support as well. Among the novel clades, NC11 was obtained with full support; however, while NC10 and NC12 were each recovered but with <50% support. Nevertheless, all three nearly-identical SSF isolates fall within previously-defined NC12 and as sister to the clade containing environmental sequences CCA33 (AY180002.1), CV1_B1_41 (AY821947.1), DB-2305-33 (EU567291.1), and ECCA80 (AY180001.1) -- themselves forming a well-supported (98% bootstrap support) clade. The two PG isolates with identical sequences, on the other hand, are only distantly related to the SSF lineage, and form a long branch with unresolved placement within “endomyxa” (*i*.*e*. are placed outside Filosa and Retaria) -- here as (unsupported) sisters to the poorly supported clade containing phytomyxids and vampyrellids. A marine anoxic environmental clade CCA44 (AY180003.1) falls within NC10 but is likely an artefact.

**Figure 3.**
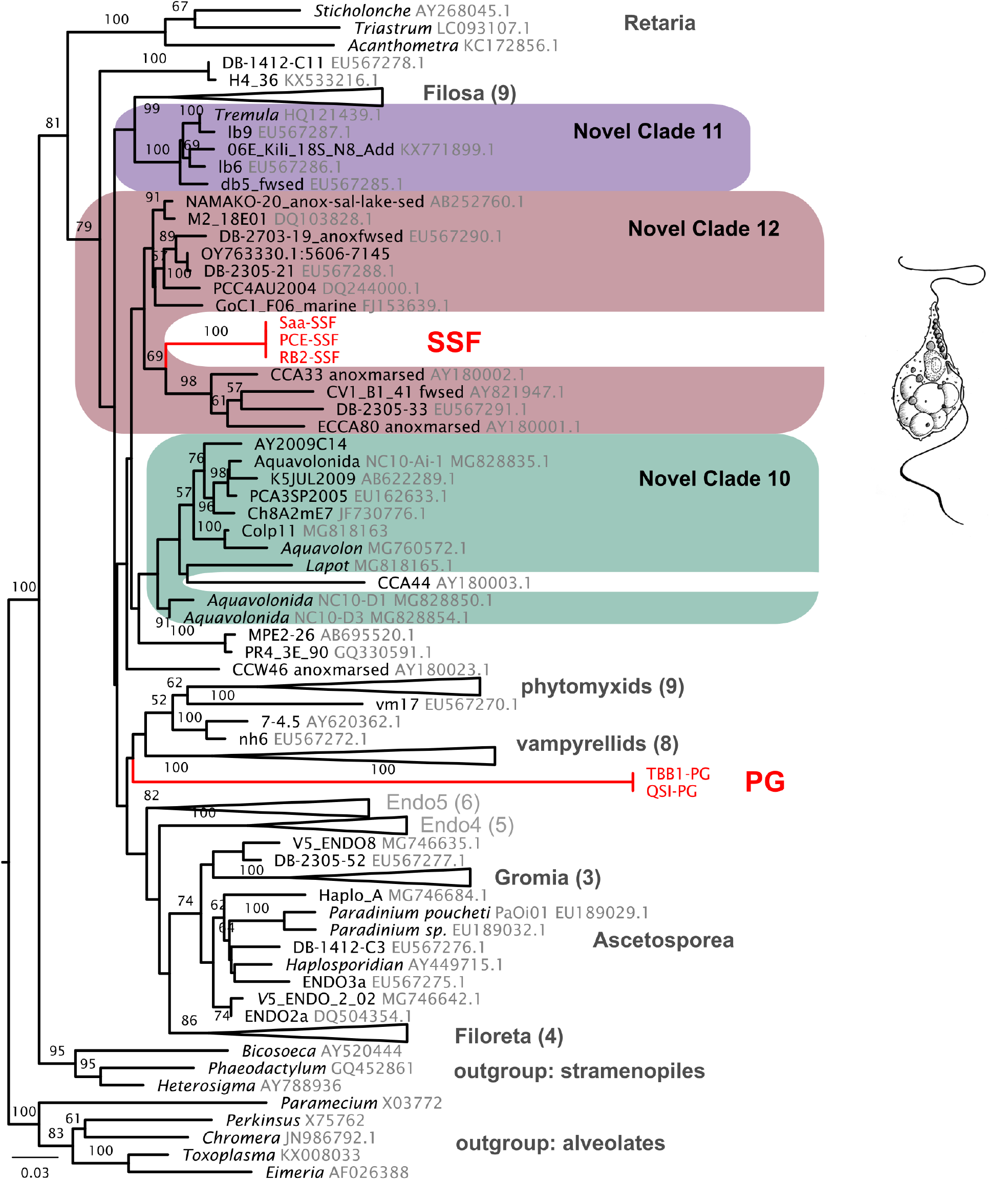
Maximum likelihood Rhizaria-focused phylogeny inferred under the GTR+Γ+I model in IQ-TREE for 1298 sites across 105 taxa with non-parametric bootstrap support values. Support values under 50% omitted. Numbers in brackets indicated number of taxa in collapsed clade.

## Discussion

We established cultures of two novel rhizarian anaerobic flagellate lineages from marine anoxic sediments, each representing a distinct adaptation to anoxia event. SSF, is the first morphologically-characterised representative of previously identified (largely from marine anoxic sediments) environmental NC12 (Bass et al., 2009). PG, on the other hand, represents a previously undiscovered deep-branching rhizarian lineage. Both organisms are presumed anaerobes and have documented eukaryotrophy on breviates.

### SSF

An organism that is certainly SSF was previously observed under light microscopy in a marine sediment sample, and described as *Cercomonas granulatus* (Lee and Patterson, 2000). Like our SSF isolates, “*Cercomonas granulatus*” was also observed to often have flagella doubled in parallel (Fig 1F). While superficially, the pliable cell shape, snake-like manner of swimming, and production of cytoplasmic strands resembles the plasticity of cercomonads (Filosa, Rhizaria), SSF and “*Cercomonas granulatus*” have a distinctive well-defined shape; features like the rostrum or row of refractile globules are absent in cercomonads. Furthermore, cercomonads are generally bacterivores or detritivores, and not eukaryotrophs. SSU rRNA gene phylogenies confirm the identity of SSF, and therefore “*Cercomonas granulatus*”, being outside cercomonads and Filosa in general. Incidentally, the “*Cercomonas* sp.” in Fig 21 O-P of Lee and Patterson (2000) is almost certainly a breviate similar to PCE-brev and Saa-brev, and likely its prey. That it was observed in late cultures (enrichments) (Lee and Patterson, 2000) is consistent with it being both an anaerobe and a eukaryotroph: presumably, anoxic conditions must develop for breviates to bloom, to then be followed by a bloom of their predator.

While it is indeed tempting to infer the row of refractile globules to be involved in active predation, we consider it unlikely for the following reasons: the globules themselves are irregular in size, which would be inconsistent with typical extrusomes; and the prey appear to respond to contact with any part of the cell surface, including the posterior end far from the globules. Thus, we propose the rows of globules are likely (e.g. lipid) storage bodies, while extrusomes involved in hunting are the small particles on the surface.

The tendency for cells to be active in presumed early division (represented by the flagellar apparatus doubled in parallel) may be reminiscent of the permanently doubled flagellar apparatus (albeit with a barely-present anterior flagellum) in the heterotrophic flagellate *Cholamonas* isolated from fly intestine (Flavin et al., 2000). However, *Cholamonas* is placed within the Sainouroidea (Cavalier-Smith et al., 2008), a filosan lineage.

### PG

Cells of PG are less structurally defined than those of SSF. PG is generally pyriform with a prominent cytoplasmic “tail” trailing up to several cell lengths. Most of the trailing flagellum remains attached to the substrate as the cell moves forward. These cells lack the row of refractile globules of SSF. While the two isolates of PG were morphologically indistinguishable, they could not be grown on each other’s prey breviates. While one cannot rule out incompatibilities arising from the background prokaryotic communities, it is also possible each of the isolates has a distinct prey preference. To our knowledge, this species has not yet been reported.

### Predation behaviour

We extensively documented both SSF and PG killing and phagocytosing the breviates and neither organism survived in culture in their absense: thus they are almost certainly obligate eukaryotrophs. Under observed conditions, neither SSF nor PG immediately ingested the prey following initial contact. In the case of PG, the prey breviate would conspicuously disintegrate within 10-20 seconds, whereas the effect appeared to be substantially slowed for SSF, taking dozens of minutes. We suspect both predators affect their prey with a toxin—presumably delivered by extrusomes—but in that case the toxin in SSF would be slower acting. In any case, prey incapacitation is decoupled from phagocytosis during the predation process, and we have observed that sometimes the prey is ultimately phagocytosed by a different individual than that which initially contacted it.

### Significance of anaerobic eukaryotrophs

Both SSF and PG are presumed to be anaerobes, based on their reliance upon an anaerobic flagellate as food source and apparently sensitivity to oxygen exposure. As SSF and PG are confidently placed apart from each other in SSU rRNA gene phylogenies, they can each be inferred as an independent anaerobic lineage, thereby providing opportunity to study two novel anaerobic metabolism systems. Prior to this study, there were two confirmed lineages with MROs among Rhizaria: *Brevimastigomonas* (Gawryluk et al., 2016) and the mikrocytids (Burki et al., 2013; Onuţ-Brännström et al., 2023)—arguably three including the denitrifying foraminiferans (Woehle et al., 2018), depending on the state of their mitochondrial reduction. In any case, we nearly double the number of known rhizarian anaerobes. It is likely more rhizarian diversity hitherto uncharacterised (or identified only through environmental metabarcoding data) lurks in anoxic environments. Having been cultivated, SSF and PG would be excellent candidates for using transcriptomics and genomics to predict anaerobic metabolic mechanisms associated with their presumed MROs.

Eukaryotrophs have been recognised as an underexplored source of novel eukaryote diversity (Tikhonenkov, 2020). They have certainly recently yielded several major eukaryotic lineages (Gawryluk et al., 2019; Janouškovec et al., 2017; Lax et al., 2018; Tikhonenkov et al., 2022). Eukaryotrophs have likewise contributed to novel rhizarian lineages of notable importance. *Aquavolon* (Bass et al., 2018) and *Lapot* (Irwin et al., 2019) both represent NC10. These findings suggest that eukaryotrophs remain an undersampled and therefore productive niche to explore for further eukaryote diversity.

Anaerobic eukaryotrophs are particularly understudied, and it is difficult to even begin exploring their role in anoxic microbial ecosystems. With the exception of ciliates and amoebae, eukaryotrophy by flagellates has been proposed to be improbable under anoxic conditions due to bioenergetic reasons (Fenchel, 2012; Muñoz-Gómez, 2023). Here, we present evidence for eukaryotrophy by eukaryotes under anoxic conditions.

### Hidden diversity lurking in environmental clades

A major portion of rhizarian diversity outside Filosa and Retaria, or “endomyxans”, is represented by environmental clades, or taxa detected only through environmental sequencing means like amplicon-based metabarcoding, or (still rarely for Rhizaria), metagenomics. Some of these environmental lineages were detected decades ago (Stoeck et al., 2003; Takishita et al., 2007, 2005) and still lack a representative with known morphology, let alone an established culture thereof. Nevertheless, two of the three “Novel Clades” relevant to this discussion, NC10-12 (Bass et al., 2009) have been characterised (discussed above). NC12, sampled almost entirely from anoxic sediments, predominantly marine, was uncharacterised until now (by SSF). It is plausible other members of the clade may also be anaerobes and/or eukaryotrophs. However, it is important to note that these clades may themselves be incredibly diverse, and the first characterised member is not always the most representative.

On the other hand, PG does not appear to match a previously defined environmental clade of rhizarians. It is not entirely surprisingly, as environmental sequencing methods are challenged by technical difficulties like primer incompatibilities. Worse yet, sediments—rich in yet-uncharacterised diversity (Cordier et al., 2022)—have been exceptionally undersampled (Vaulot et al., 2022), which likely presents a problem to detection of diversity particularly among endomyxa (Berney et al., 2022). Furthermore, both anaerobes or eukaryotrophs have been traditionally difficult to culture and work with. Combination of cultivation-independent approaches, such as single-cell transcriptomics (Kolisko et al., 2014) with classical cultivation of organisms of interest identified by light microscopy could at least improve reference databases for classifying existing metabarcoding data. The prime environments to begin this search in would include anoxic sediments, with particular focus on the anaerobic eukaryotrophs previously considered to be unlikely. Altogether, these novel rhizarian lineages, particularly among the “endomyxans”, would enable us to finally better resolve rhizarian phylogeny and, perhaps, locate its root.

## Acknowledgements

We would like to thank Claudio Slamovits (Dalhousie University) and Susana Breglia (Dalhousie University) for supplying the sample for TBB1-PG. Gordon Lax (Dalhousie University and University of British Columbia) for supplying the sample for QSI-PG. Greg Gavelis (University of British Columbia and Bigelow Labs) provided the Saa-SSF (and Saa-brev) sample, while Noèlia Carrasco (IRTA) provided sampling access and the permit for the source of LRM1b, upon which TBB1-PG was maintained. We also thank David Bass (Cefas) for discussion on rhizarian novel clades. The research was supported by NSERC grant RGPIN-2019-04336 to RMRG and 298366-2014 (&-2019) to AGBS.

