## Supplemental Table 1. for "Two newly cultivated eukaryotrophic flagellates represent distinct anaerobic lineages within Rhizaria"

### Tables

| Isolate ID | Location name | Sample type | Sample notes | Latitude | Longitude | Salinity (ppt) | Sample credits | Date |
|---|---|---|---|---|---|---|---|---|
| **PCE-SSF** | PEI Cavendish Eelgrass | subtidal sediment | anoxic | 46°30'00.0"N | 63°25'11.3"W | marine | | July, 2016 |
| **Saa-SSF** | Centennial Beach of White Rock, BC | mud at low tide | top cm | 49°00'N | 123°01'W | marine | Greg Gavelis | May, 2015 |
| **RB2-SSF** | Roberts Bay, Sidney, BC | mud at low tide | anoxic | 48°66"N | 123°40"W | marine | | August ,2023 |
| **TBB1-PG** | Tsawwassen (Boundary Bay) | Intertidal mud flat | anoxic | 49°00'27.1"N | 123°01'52.7"W | marine | Claudio Slamovits | May, 2015 |
| **QSI-PG** | Small Inlet, Quadra Island, BC | marine sediment | rich in shellfish | 50°15'43.1"N | 125°16'17.5"W | marine | Gordon Lax | 2016 |

**Table 1**. Details on sampling locations for the isolates discussed in this paper.
