## Supplemental Table 2. for "Two newly cultivated eukaryotrophic flagellates represent distinct anaerobic lineages within Rhizaria"

| Isolate ID | Fwd Primer | Rev Primer | Temp (°C) |
| --- | --- | --- | --- |
| PCE-SSF | 25F | 1498R | 55 |
| Saa-SSF | 82F | 1498R | 60 |
| RB2-SSF | SSF18S_LrgF3 | SSF18S_LrgR3 | 53 |
| TBB1-PG | EukA | EukB | 53 |
| QSI-PG | EukA | EukB | 54 |

**Table 2**. PCR conditions. Details on primers used to amplify the SSU rRNA gene for the isolates described here. Temp refers to the annealing temperature used in PCR amplification.
