## Supplemental Table 3. for "Two newly cultivated eukaryotrophic flagellates represent distinct anaerobic lineages within Rhizaria"

| Name | Fwd or Rev | Sequence (5′-3′) |
|---|---|---|
| EukA | Fwd | AACCTGGTTGATCCTGCCAGT |
| EukB | Rev | TGATCCTTCTGCAGGTTCACCTAC |
| 25F | Fwd | CATATGCTTGTCTCAAAGATTAAGCCA |
| 82F | Fwd | GAAACTGCGAATGGCTC |
| SSF1345R | Rev | TAATCTAGCCCCATCACGTTGCA |
| 1492R | Rev | AAGTCGTAACAAGGT |
| 1498R | Rev | CACCTACGGAAACCTTGTTA |
| SSF18S_LrgF3 | Fwd | TAATCTAGCCCCATCACGTTGCA |
| SSF18S_LrgR3 | Rev | ACGAAACCCAATTCAGCATCA |

**Table 3**. Primer sequences. Sequences of the primers used to amplify the SSU rRNA gene in this study, as reported in Table 2.
